# Direct observation of aggregate-triggered selective autophagy

**DOI:** 10.1101/2021.04.21.440799

**Authors:** Anne F.J. Janssen, Giel Korsten, Wilco Nijenhuis, Eugene A. Katrukha, Lukas C. Kapitein

## Abstract

Degradation of aggregates by selective autophagy is important as damaged proteins may impose a threat to cellular homeostasis. Although the core components of the autophagy machinery are well-characterized, the spatiotemporal regulation of many selective autophagy processes, including aggrephagy, remains largely unexplored. Furthermore, because most live-cell imaging studies have so far focused on starvation-induced autophagy, little is known about the dynamics of aggrephagy. Here, we describe the development and application of the mKeima-PIM assay, which enables live-cell observation of autophagic turnover and degradation of inducible protein aggregates in conjunction with key autophagy players. This allowed us to quantify the relative timing and duration of different steps of aggrephagy and revealed the short-lived nature of the autophagosome. The assay furthermore showed the spatial distribution of omegasome formation, highlighting that autophagy initiation is directly instructed by the cargo. Moreover, we found that nascent autophagosomes mostly remain immobile until acidification occurs. Thus, our assay provides new insights into the spatiotemporal regulation and dynamics of aggrephagy.

## INTRODUCTION

Efficient turnover of misfolded proteins and damaged or redundant organelles is essential to maintain cellular homeostasis and cells have evolved multiple pathways to ensure (protein) quality control. Misfolded proteins can be refolded by chaperone networks or degraded via the ubiquitin-proteasome system or the autophagy-lysosome pathway. Superfluous organelles can be degraded as a whole by macroautophagy (autophagy hereafter). Although autophagy was initially characterized as a bulk degradation pathway, it has become increasingly clear that it serves important roles in the selective degradation of cytoplasmic material in order to maintain homeostasis (Anding and Baehrecke, 2017; Kirkin and Rogov, 2019). Bulk autophagy is induced by nutrient deprivation and serves to replenish essential metabolites, whereas in selective autophagy substrates or cargos, such as damaged mitochondria, intracellular pathogens or aggregates, are specifically degraded to avoid the possible danger these may impose on cellular homeostasis. The importance of selective removal of damaged proteins through autophagy (aggrephagy) has become clear with the identification of mutations in autophagy receptors that cause neurodegenerative disease associated with protein aggregation (Deng et al., 2017; Rui et al., 2015).More knowledge on aggrephagy might enable the design of strategies that interfere with this specific autophagic processes and lead to novel therapies.

In autophagy a double-membrane vesicle, the autophagosome, forms around cytoplasmic cargo. First a crescent-shaped cisterna, the isolation membrane, is formed within omegasomes, ER subdomains enriched in phosphatoidyl-3-phosphate (PI(3)P) (Axe et al., 2008). The isolation membrane is subsequently elongated and closed to complete autophagosome formation, after which it will fuse with lysosomes to ensure degradation. Although the core components of autophagy are well-known, the exact timing and spatial regulation of selective autophagy processes are not well-described. Starvation-induced autophagosomes form throughout the cell and then concentrate in the perinuclear area, where they fuse with lysosomes (Jäger et al., 2004; Jahreiss et al., 2008). Much less is known about the spatiotemporal dynamics of aggrephagy. For example, it is not known whether autophagosomes directly form at the site of aggregate formation, how long mature autophagosome containing an aggregate exist before fusion with lysosomes, and where this fusion occurs.

Answering these questions requires direct observation of aggrephagy. Previous work has focused on imaging of starvation-based autophagy (Axe et al., 2008; Itakura and Mizushima, 2010; Jahreiss et al., 2008; Karanasios et al., 2016; Koyama-Honda et al., 2013; Tsuboyama et al., 2016), mitophagy (Dalle Pezze et al., 2020; Zachari et al., 2019) and xenophagy (Kageyama et al., 2011). While this has revealed the timing of recruitment of specific autophagy proteins, it has remained unclear to which extent these findings apply to aggrephagy. Moreover, in these previous studies it remained unexplored how the successive recruitment of autophagy mediators relates to successful cargo degradation.

Recently, we introduced the PIM assay as an inducible assay to study aggrephagy (Janssen et al., 2018). The PIM assay allows for the inducible formation of proteinaceous clusters inside living cells by rapalog2-induced multimerization of the PIM protein, which is comprised of several homodimerization domains. We have previously shown that these clusters behave as aggregates inside the cell and are targeted to the lysosome via the selective autophagy pathway (Janssen et al., 2018). This assay uniquely allows the direct observation of autophagic aggregate clearance as well as its dynamics and spatio-temporal regulation. We previously employed the dual EGFP-mCherry tag to visualize transfer of aggregates to the lysosome by the selective loss of EGFP fluorescence from the PIM clusters. However, this approach requires two optimal channels for live-cell imaging and therefore precludes straightforward labelling and high-contrast imaging of other proteins in conjunction with autophagic turnover of aggregates. Preferably, the GFP channel would be left available to facilitate the imaging of such proteins at low expression levels. Here we describe the development of the mKeima-PIM assay, which uses a single-color pH-sensitive fluorophore to enable live-cell imaging of aggrephagy in combination with key markers of various steps in the autophagy pathway, such as the PI(3)P-binding protein DFCP1, the autophagosomal SNARE protein STX17, and the late endosomal/lysosomal marker RAB7. This enabled us to unravel the spatiotemporal dynamics of the autophagic clearance of aggregates.

## RESULTS

To combine direct observation of aggrephagy with imaging specific players, we decided to exchange the dual tag in the PIM construct for the mKeima fluorophore (Figure 1a). mKeima is a pH-sensitive fluorophore whose unique spectral properties makes it very useful for the autophagy field (An and Harper, 2018; Katayama et al., 2011; Lazarou et al., 2015). Specifically, mKeima emission peaks at 620 nm and, importantly, has a bimodal excitation spectrum with peaks at 440 and 568 nm. While the emission profile remains unchanged, the efficiency of excitation is pH-sensitive and shifts from 440 nm at neutral pH (cytoplasmic) to 568 nm for acidic pH (lysosome). Moreover, mKeima is, like mCherry, resistant to lysosomal proteases and imaging can be combined with green-emitting fluorophores (Katayama et al., 2011).

**FIGURE 1:**
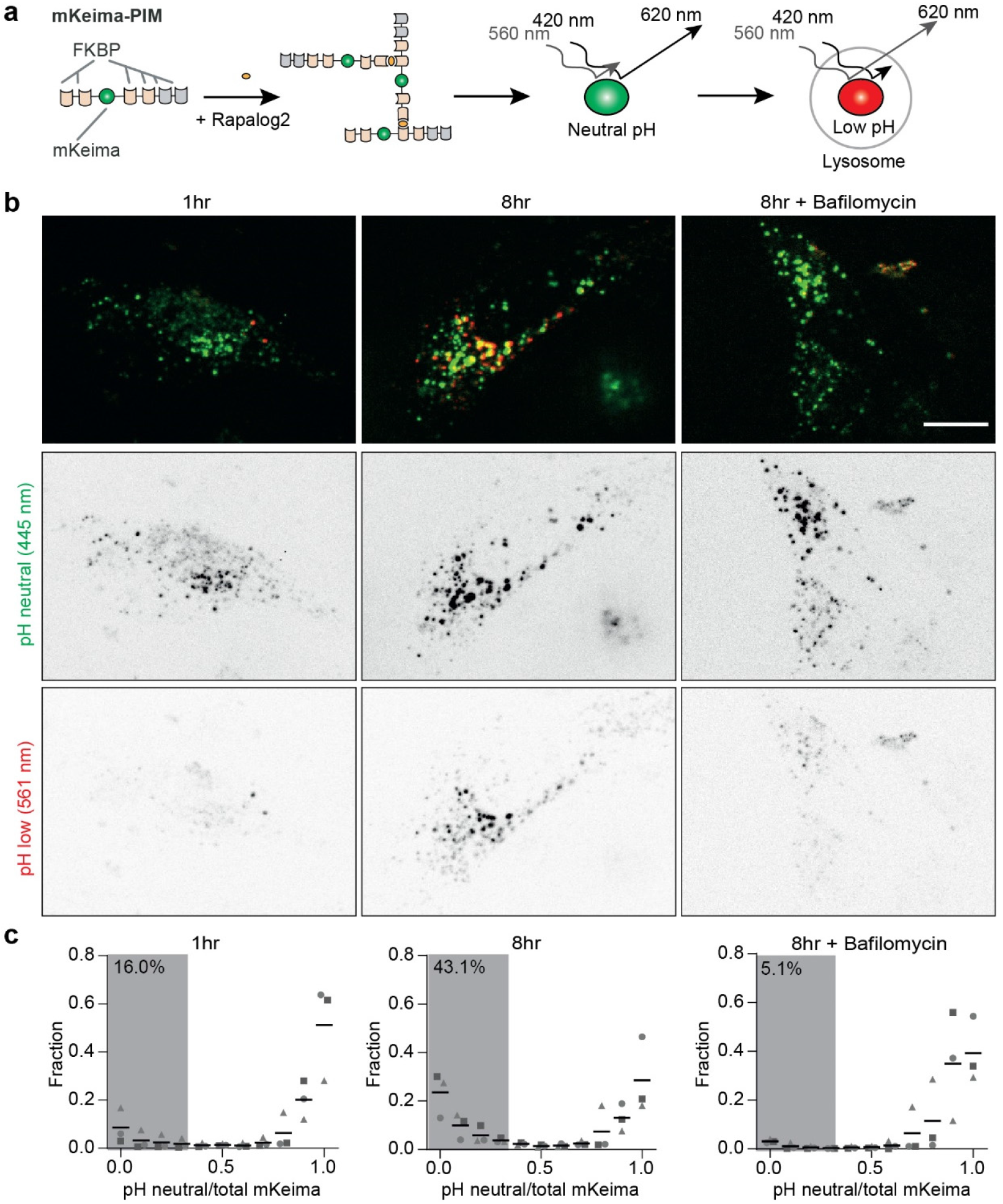
The mKeima-PIM assay enables following aggrephagy using a single emission channel. a) Assay: mKeima-PIM aggregates are formed by multimerization upon rapalog2 addition. Peak emission of mKeima fluorophore is at 620 nm, but the excitation spectrum depends on pH. Aggregates are pseudo-coloured throughout this paper: green for excitation at ∼420 nm and red for excitation at ∼560 nm. b) HeLa cells expressing mKeima-PIM at different timepoints after cluster formation by addition of rapalog2. Inverted contrast grayscale panels show mKeima emission at neutral pH by using 445 nm excitation (middle row) and emission of mKeima at low pH with 561 nm excitation (bottom row). Cells in third column were treated with 200 nM Bafilomycin A1. c) Distribution of the ratio of mKeima fluorescence intensity, defined as I_mKeima-neutral pH_/I_total mKeima_ at 1 h (left) and 8 h (middle) after cluster formation, and 8 h after formation in the presence of Bafilomycin. Each dataset contains 500-1600 clusters and 3 independent experiments were analysed (indicated by circles, triangles and squares). The mean from 3 independent experiments is indicated. The percentage of clusters in low pH environments is indicated as the average fraction of clusters with a ratio <0.35 of 3 independent experiments. Scale bar, 10 µm.

First, we expressed mKeima-PIM in HeLa cells to test whether the replacement of the dual EGFP-mCherry tag with mKeima still enabled the formation and clearance of aggregates. Indeed, we observed clear aggregate formation 1 h after rapalog2 addition (Figure 1b). When we subsequently performed dual excitation ratiometric imaging of mKeima, we observed that the efficiency of mKeima excitation with blue light (445 nm) was reduced in favour of excitation with yellow light (561 nm) for a subpopulation of aggregates at 8 h after aggregate formation. This shift in excitation sensitivity was largely prevented by treatment with Bafilomycin A1, which is a vacuolar-type H^+^-ATPase inhibitor (Yoshimori et al., 1991) and blocks autophagosome-lysosome fusion and lysosomal degradation (Yamamoto et al., 1998). Our results therefore show that identically to the dual EGFP-mCherry-PIM aggregates, the mKeima-PIM aggregates are cleared by autophagy.

To quantify the clearance of aggregates by autophagy we performed automated puncta detection using the ComDet V.0.4.1 plugin for ImageJ. This plugin automatically detects puncta and subsequently quantifies fluorescence in the detected area in both channels. From these values we get the mKeima ratio, defined as the mKeima fluorescent intensity after 440 nm excitation divided by the total mKeima emission (440 and 568 nm excitation). This ratio was calculated for each particle and plotted in a histogram as a fraction of particles (Figure 1c). A ratio close to one indicates that mKeima is mostly sensitive to blue light excitation and that the aggregate resides in a neutral environment, while a low ratio indicates an acidic environment, i.e. the lysosome. When comparing samples treated with rapalog2 after 1 h or 8 h, we indeed observed an increase in the number of particles with low ratios 8 h after aggregate formation (16.0% versus 43.1%), which was largely prevented by treatment with Bafilomycin A1 (5.1%).

To examine whether the mKeima-PIM assay would also be compatible with high throughput approaches, we next measured mKeima intensity with both excitation channels in a population of cells by FACS (Figure 2). Here, 20.5 % of cells displayed a spectral shift at 8 h after rapalog2 addition, compared to 0.69 % at 1 h after rapalog2 addition, and this shift was completely abolished upon treatment with Bafilomycin A1. Thus, mKeima-PIM enables robust and high-throughput detection of autophagic flux following aggregate induction.

**FIGURE 2:**
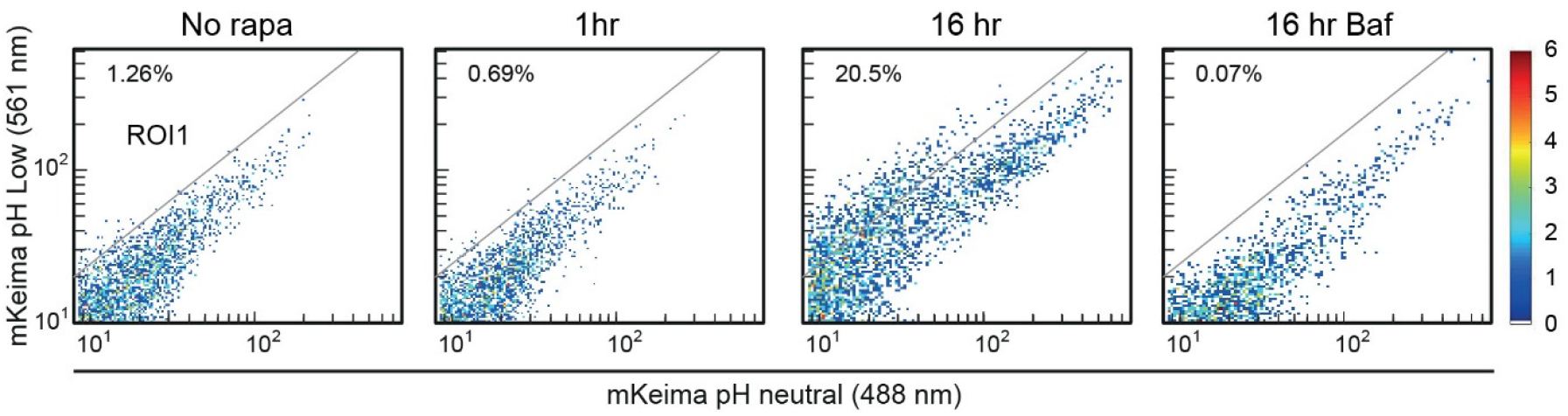
FACS-based analysis of aggrephagy progression using mKeima-PIM. mKeima-PIM clearance followed by flow cytometry. HeLa cells expressing mKeima-PIM were treated with rapalog2 and analysed by FACS for mKeima-PIM at low and neutral pH. Cells in the right panel were treated with Bafilomycin A1. Percentage of cells in ROI1 is indicated.

Next, we aimed to directly study autophagosome initiation with respect to cargo in aggrephagy and we therefore generated a Double FYVE-Containing Protein-1 (DFCP1)-GFP stable cell line (Figure 3a). DFCP1 is a PI(3)P-binding protein that localizes to omegasomes at the onset of autophagosome formation (Axe et al., 2008). During live-cell imaging at least 4 h after PIM formation, we indeed frequently observed a DFCP1 positive signal appearing at the site of an aggregate (Figure 3b,c). Simultaneous ratiometric imaging of mKeima revealed that, in all cases, DFCP1 emerged and disappeared at the PIM site before acidification. DFCP1 disappeared from the aggregate 5 - 15 minutes before the onset of acidification (Figure 3e). In several cases, DFCP1 was forming the ring-like structures observed previously for starvation-induced autophagy (Figure 3d), but we typically observed smaller puncta (Figure 3b-c). These small puncta could potentially still represent ring-like structures that could not be resolved by diffraction-limited microscopy. Interestingly, most aggregates were largely immobile before and during the presence of the DFCP1 signal (Figure 1d), but became increasingly mobile after loss of DFCP1 from the aggregate, which coincides with the release of newly formed autophagomes from the omegasome (Figure 3f). These observations suggest that aggregates might be tethered to a potential autophagosome formation site before and during autophagosome formation.

**FIGURE 3:**
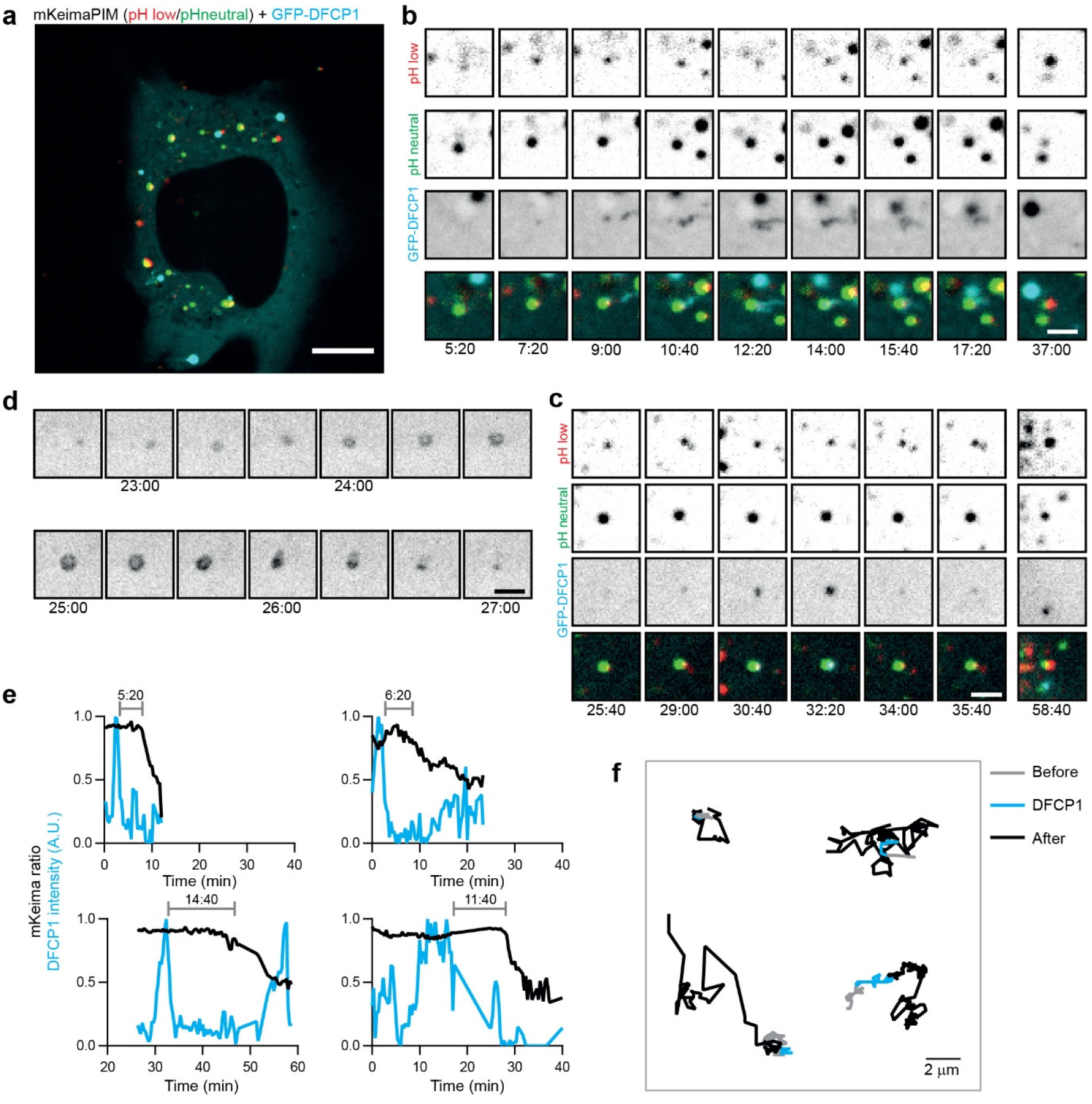
Local omegasome formation at aggregates precedes acidification by around 7 minutes. a) HeLa cell stably expressing GFP-DFCP1(Cyan) and transiently expressing mKeima-PIM (red and green for neutral and low pH, respectively). b-c) Zoom of an aggregate acquiring DFCP1 signal before acidification. d) Time lapse imaging of ring structure formation by DFCP1. e) Time lapse analysis of DFCP1 intensity and mKeima ratio of individual aggregates, showing that DFCP1 is acquired before acidification. The time between presence of DFCP1 and start of mKeima-PIM acidification is indicated. f) Analysis of mobility of aggregates before (gray), during(cyan) and after (black) DFCP1 signal was present around the aggregate. Scale bars, 10 µm (a) and 2 µm (b,c,d).

To visualize completion of autophagosome formation, we generated a stable GFP-STX17 cell line. STX17 is the autophagosomal SNARE involved in fusion of the autophagosome with late endosome (LE)/lysosome (Itakura et al., 2012), and is recruited at the end of autophagosome formation, probably by mATG8s (Kumar et al., 2018). STX17 is not recruited to the isolation membrane and is therefore a useful marker to identify fully formed autophagosomes (Itakura et al., 2012). In line with previous observations (Itakura et al., 2012), we observed STX17 localization to the ER and mitochondria under normal nutritional conditions (Figure 4a-b). In addition, we observed punctate (arrowheads) and ring-shaped (arrow) structures that suggested the presence of autophagosomes >4 h after aggregate induction.

**FIGURE 4:**
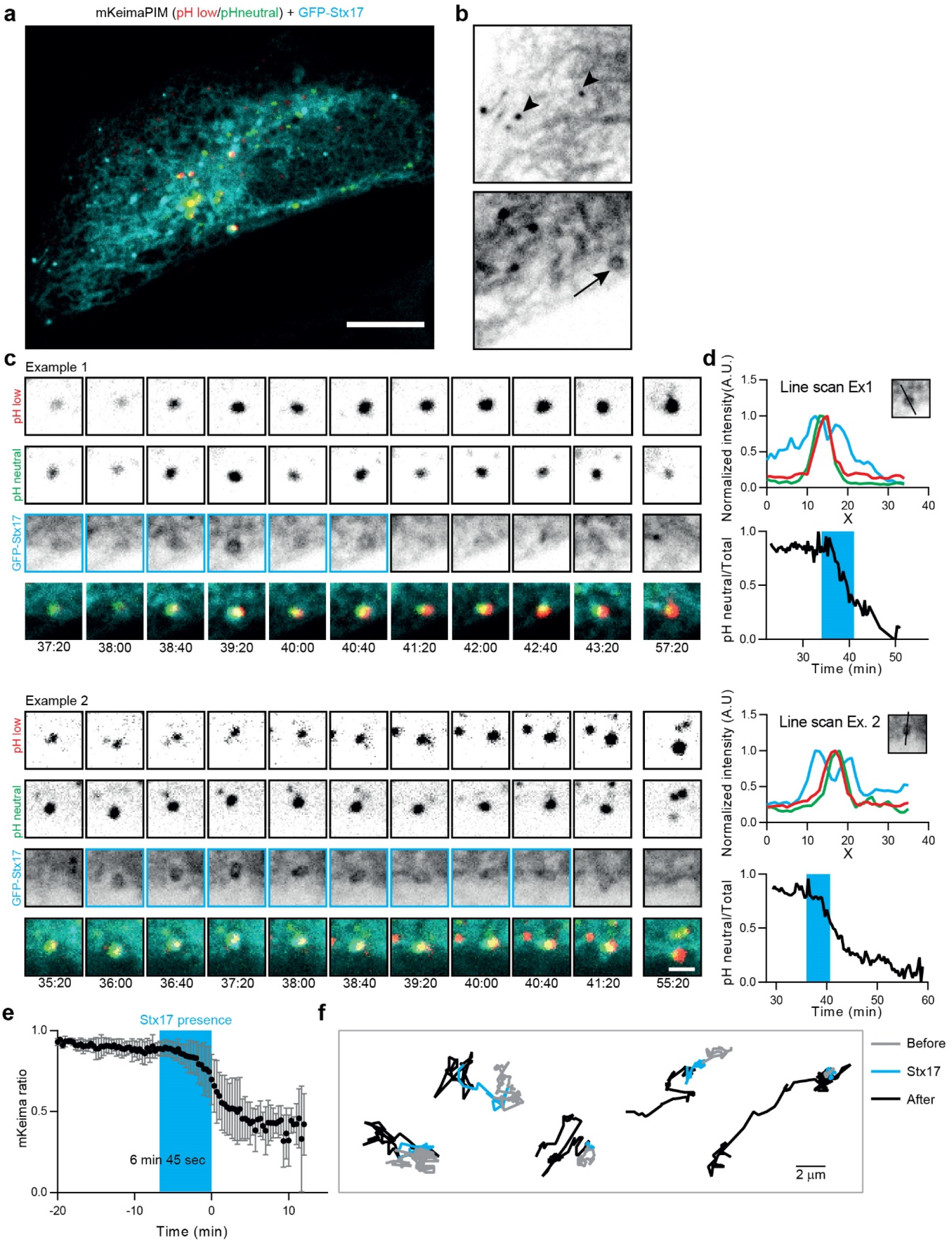
Local autophagosome formation at aggregates is directly followed by acidification. a) HeLa cell stably expressing GFP-STX17 and transiently expressing mKeima-PIM b) Zoom of cell in (a) showing GFP-STX17 localization to ER and mitochondria and the presence of small vesicles (arrowhead) and rings (arrows) positive for STX17. c) Two examples with selected frames from a time-lapse movie showing STX17 recruitment to the aggregate and subsequent loss of STX17 concomitant with acidification. d) Analysis of time-lapse examples in (c) showing intensity profiles indicating presence of the aggregate inside a STX17 positive autophagosome (top) and the mKeima ratio over time (bottom). The blue box indicates when a STX17 positive signal at the aggregate was observed. e) Average mKeima ratio over time and indication of average STX17 signal presence. Data represents mean ± s.d. from 12 aggregates. Events were aligned according to the first frame after STX17 disappearance. f) Analysis of mobility of aggregates before, during and after STX17 signal was present around the aggregate. Scale bars, 10 µm (a) and 2 µm (c).

During time-lapse imaging of preformed aggregates (4-8 h after rapalog2 addition), we clearly observed the formation of STX17-positive rings around aggregates just before acidification of the cargo (Figure 4c-d and Supplementary Video 1). Following the onset of acidification, STX17 was released from the aggregate-containing autophagosomes (Figure 4d). On average, STX17 was associated with aggregates for about 7 min. At the time of STX17 disappearance, the acidification ratio had proceeded to about 30% of the total change (Figure 4e). This suggests that once fully formed, autophagosomes quickly fuse with LE/Lysosomes. Moreover, by monitoring the displacements before, during and after STX17 association, we found that aggregates remain largely immobile before and during the presence of STX17 (Figure 4f). Only after STX17 disappearance, marking the fusion with LE or lysosome, motility is increased. While the conventional view holds that autophagosomes are transported to lysosomes in order to form autolysosomes, these results demonstrate that in aggrephagy the reverse scenario is more likely. The nascent autophagosomes remain immobile and potentially tethered until acidification, when amphisome or autolysosome formation has occurred through fusion with LE or lysosomes, respectively.

When expressed at higher levels, bigger aggregates can form with more amorphous structures than the largely spherical aggregates we described so far. To see whether autophagosomes could also form around these bigger structures, we imaged STX17 together with these larger mKeima-PIM aggregates (Figure 5a). Interestingly, while we did observe the association of STX17 with these big aggregates (Figure 5b), there was no clear indication that STX17 release coincided with autophagosome maturation and subsequent degradation. The dissociation of STX17 from the aggregate is believed to indicate successful breakdown of the inner autophagosomal membrane (IAM) (Tsuboyama et al., 2016) and thereby the onset of degradation and acidification. Nonetheless, we did not observe any mKeima shift in these aggregates, which raises the question how and why STX17 leaves these autophagosomal structures. One possibility would be that STX17 was erroneously recruited before autophagosome closure, which has been observed previously (Tsuboyama et al., 2016). This is supported by our observation that also for smaller aggregates, not every STX17 recruitment event led to degradation. In several cases, we observe either the loss of STX17 without concomitant acidification (Figure 5c-d) or the stable presence of STX17 signal on aggregates (Figure 5e-f). In the latter case, STX17 remained localized to the aggregate even though acidification of the autophagosome had already started, based on the decrease in the mKeima ratio. Based on the earlier finding that IAM breakdown, rather than acidification, causes STX17 dissociation(Tsuboyama et al., 2016), this suggests that IAM breakdown is delayed or blocked.

**FIGURE 5:**
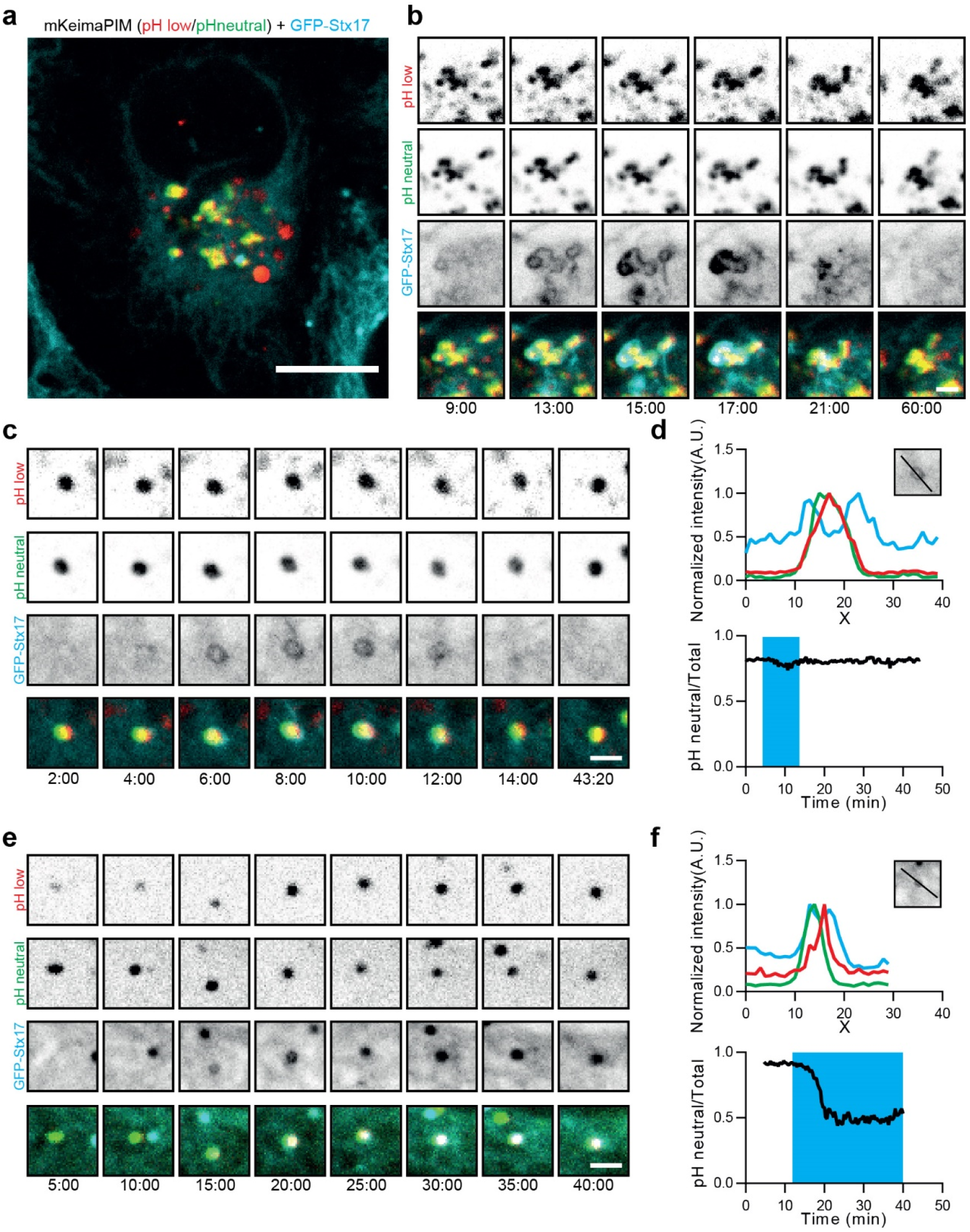
STX17 recruitment does not always result in successful clearance. a) HeLa cell stably expressing GFP-STX17 and transiently expressing mKeima-PIM. b) Selected frames from a time-lapse movie showing STX17 recruitment and dissociation from a larger aggregate not followed by acidification. c) Example showing frames from a time-lapse movie where STX17 recruitment did not lead to subsequent acidification. d) Analysis of the time-lapse examples in (c) showing the presence of the aggregate inside the STX17 positive autophagosome and the mKeima ratio over time. The blue box indicates the time where a STX17 positive signal at the aggregate was observed. e) Frames from a time-lapse movie showing prolonged recruitment of STX17 leading to partial but not full acidification. f) Analysis of example shown in (e) with an intensity profile and the mKeima ratio over time. The blue box indicates the time where a STX17 positive signal at the aggregate was observed. Scale bars, 10 µm (a) and 2 µm (b,c,e).

The final steps of autophagy involve multiple fusion and kiss-and-run events with endosomes and lysosomes to enable acidification and final degradation of the autophagosomal content (Dunn Jr., 1990; Jahreiss et al., 2008; Zhao and Zhang, 2018). RAB7 is a member of the Rab family of small GTPases and localizes to LE and lysosome (Guerra and Bucci, 2016; Jimenez‐Orgaz et al., 2018).To directly examine the interplay between aggregate-containing autophagosomes and both LE and lysosomes, we tagged endogenous RAB7 with GFP using CRISPR-Cas9-mediated genome editing (Supplementary Figure 1a-b). The generated GFP-RAB7 cell line clearly shows vesicular structures ranging in size with bigger vesicles close to the nucleus and smaller vesicles at the periphery (Figure 6a-b). Aggregates with low mKeima ratios (low pH, red arrows) were typically associated with higher RAB7 levels than aggregates with high mKeima ratios (high pH, green arrows; Figure 6c). When we monitored an aggregate over time, we often observed multiple contacts with RAB7 positive vesicles and a gradual increase in RAB7 signal together with a decrease in mKeima ratio (Figure 6d,e). RAB7 positive vesicles that contacted the forming autolysosome would sometimes fuse (Figure 6g and Supplementary Video 2), but most often left after several seconds, reflecting potential kiss-and-run events to transfer membrane-bound proteins and/or proteolytic enzymes (Figure 6f). In perinuclear regions crowded with RAB7 positive vesicles, forming autolysosomes showed multiple rounds of transient contact with RAB7, eventually resulting in content acidification (Supplementary Figure 2a-b and Supplementary Video 3). These observations demonstrate that multiple contact events and fusions are often needed for the full acidification and autophagosomal degradation of aggregates.

**FIGURE 6:**
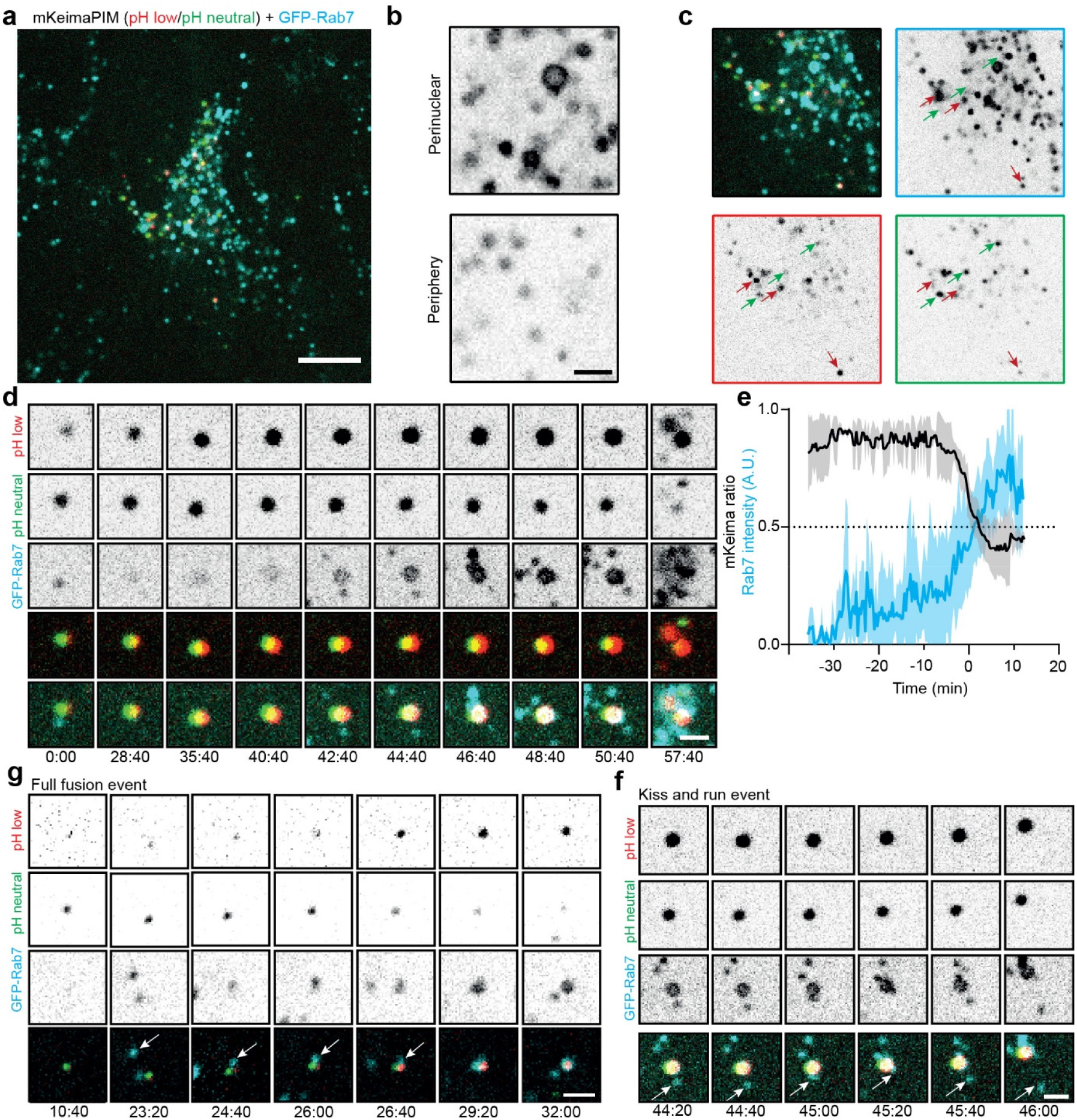
Rab7 is gradually recruited to aggregates through multiple fusion and contact events. a) Endogenous GFP-RAB7 KI HeLa cell line transiently showing mKeima-PIM aggregates. b) Endogenous GFP-RAB7 signal showing larger RAB7 positive vesicles at the perinuclear area and smaller vesicles in the periphery. c) Zoom of the cell in (a) showing aggregates at neutral pH (green) and low pH (red) and the correlation with RAB7. d) Selected frames from a time-lapse movie showing the gradual recruitment of RAB7 and concomitant decrease of mKeima ratio. e) Analysis of RAB7 intensity (cyan) and mKeima ratio (black) of a 10 typical acidification events, including the one shown in (d). Data shown are mean (solid line) ± s.d. (shaded area) of 3-10 events. f-g) Time-lapse imaging of GFP-RAB7 KI HeLa cells showing putative kiss-and-run events with RAB7 positive vesicles (g) and full fusion event of RAB7 positive vesicle with the autolysosome (f). Scale bars: 10 µm (a) and 2µm (b,d-g)

## DISCUSSION

Here we have introduced mKeima-PIMs as an inducible probe for the autophagic clearance of aggregates. Compared to the dualPIMs that we previously introduced, mKeima-PIMs offer the important advantage that they report on acidification using only the red emission channel, enabling the combined imaging with GFP-labelled proteins. This facilitates the use of cell lines with stably expressing GFP-labelled proteins and enabled us to directly visualize aggregate clearance in conjunction with different key autophagy factors. Together, our results demonstrate that during aggrephagy, the isolation membrane starts forming from the omegasome at the aggregate site about 5-15 minutes before start of cargo acidification (Figure 7). The lifetime of the omegasome itself is only a few minutes. Completion of autophagosome formation, as identified by STX17 recruitment, is in most cases directly followed by autophagosome acidification. The mature autophagosome has an average lifetime of around 7 minutes before acidification and STX17 dissociation. Finally, RAB7 gradually accumulates on the amphisome/autolysosome by multiple kiss-and-run and full fusion events, resulting in a drop in pH and final cargo degradation.

**FIGURE 7:**
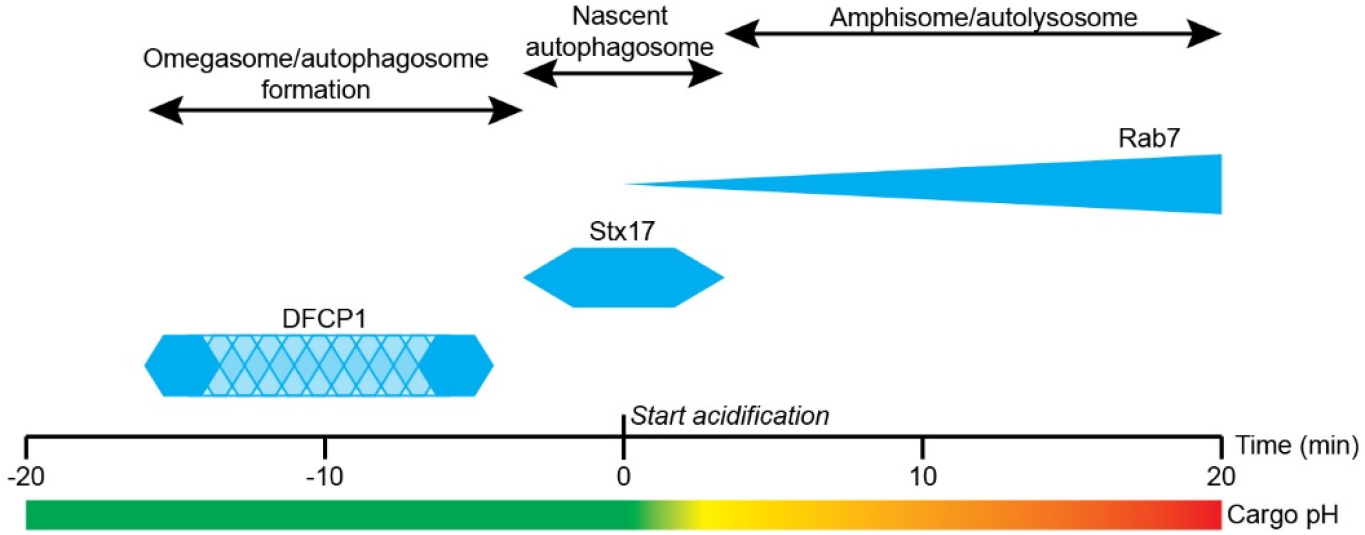
Timing of autophagy factor recruitment with respect to aggregate acidification. First, DFCP1 is recruited to aggregates (omegasome formation). DFCP1 puncta are present only for a short time but the timing of DFCP1 puncta presence relative to acidification is variable (patterned area). Just before acidification of the cargo, STX17 is recruited for ∼7 min and STX17 leaves the autophagosome after start of cargo acidification. RAB7 is gradually recruited to the autophagolysosome while cargo acidification is also taking place.

Previous work has used live-cell imaging to analyse the temporal sequence in which different autophagy-related proteins are recruited to autophagic sites following starvation (Koyama-Honda et al., 2013). This revealed that p62/SQSTM, a receptor and substrate for selective autophagy, accumulated at autophagic sites after appearance of DFCP1/ZFYVE1, suggesting that substrates for autophagy are transported to preformed omegasomes. In contrast, our results show that the omegasome (which is DFCP1-positive) is formed at the cargo site. This suggests that autophagosome formation is, at least partially, instructed by the aggregates. Indeed, cargo-instructed autophagy was recently proposed for selective mitophagy where accumulation of the autophagy receptor NDP52 on mitochondria leads to localization and activation of the ULK1 complex to induce autophagosome formation (Vargas et al., 2019). Recruitment of upstream autophagy machinery by autophagy receptors was also proposed for xenophagy (Ravenhill et al., 2019) and p62 condensates (Turco et al., 2019) and resembles the situation described for the Cvt pathway in yeast, where, unlike non-selective autophagy, the cargo is required for proper autophagosome formation (Shintani and Klionsky, 2004). Such a cargo-instructed process could facilitate the selective clearance of specific items and prevent engulfment of other cytoplasmic factors (Rogov et al., 2014; Sawa-Makarska et al., 2014; Zaffagnini and Martens, 2016).

Using our live-cell approach, we found that aggregate-containing autophagosomes are relatively short-lived structures and that fully formed autophagosomes only persist for about 7 minutes before STX17 dissociation and complete acidification of the autolysosome. This is consistent with the lifetime of STX17 positive autophagosomes in starvation conditions (Tsuboyama et al., 2016). In addition, we could demonstrate that maturation into autolysosomes/amphisomes and the corresponding acidification typically takes place at the site of autophagosome formation through multiple fusion or kiss-and-run events. Thus, in contrast to what has been shown for starvation-induced autophagy (Jahreiss et al., 2008), aggregate-triggered autophagosomes are largely immobile and depend on motile lysosomes and late endosomes for acidification near their site of formation.

In comparison to earlier work that performed live-cell imaging of autophagy, an important advantage of mKeima-PIM is that it enables simultaneous probing of cargo fate and different components of the autophagy machinery. This revealed, for example, that often the inner part of the autophagosome already starts to acidify in the presence of STX17, presumably before IAM breakdown. In addition, it revealed that recruitment and release of STX17 is not always a predictor of successful acidification.

In addition to providing new imaging opportunities, the use of mKeima was also beneficial for FACS. Using dual EGFP-mCherry PIM, we previously reported that a fraction of PIMs displayed a spectral change in the presence of Bafilomycin A1, most likely as a result of GFP self-quenching (Janssen et al., 2018). In contrast, using mKeima-PIMs we observed no spectral changes in this condition and we therefore recommend this approach for high-throughput analysis of autophagy using FACS. Nonetheless, for image-based analysis of autophagy progression we recommend using the dual EGFP-mCherry PIM in fixed samples. Robust ratiometric imaging can be hampered by aggregate displacement within the time required to take two sequential images. As such, analysing fixed samples would enable more reliable analysis. Because the pH gradient across the lysosomal membrane is lost upon fixation, mKeima cannot be used in fixed samples. In the dual EGFP-mCherry PIM assay, GFP is degraded and therefore the colour switch observed on lysosomal transfer is preserved upon fixation. The two PIM assays thus have benefits for different types of experiments, which together will likely lead to a deeper understanding of aggrephagy.

## METHODS

### Plasmids

The mKeima PIM construct was an adapted version of our PIM construct published before (Janssen et al., 2018). We exchanged the mCherry and EGFP by mKeima resulting in a construct consisting of four FKBP homodimerization domains with sequence variation, mKeima and two FKBP heterodimerization domains in the mammalian expression vector pβactin. The first two FKBP homodimerization domains contain three mutations (V24E, Y80C, and A94T) that were found to aid multimerization. mKeima was amplified by pcr using mKeima-RED-N1 (Addgene #54597) as a template. mKeima-RED-N1 was a gift from Michael Davidson.

pDONOR-GFP-RAB7 encodes a tagging module, consisting of a 6xGGGGS linker and a N-terminal EGFP tag, flanked by 1000 bp homology arms that are homologous to the genomic region immediately surrounding the RAB7 start codon and was generated by PCR and Gibson assembly strategies. Primers used to amplify the 5’ homology arm were 5’-ggagatcggtacttcgcgaatgcgtcgagatgcggccgctaccctggcaaatgagaggc-3’ and 5’-tccgttccagtgtggttgccagcatggtggcgcgccccttcaaactaaagggggaaaag-3’. Primers used to amplify the 3’ homology arm were 5’-ggagggggttctggtggtggtagctacgtaacctctaggaagaaagtgttgctg-3’ and 5’-tgcactcgtcggtcccggcatccgatacgcgtgcggccgcctgggcaaatgctagcgaac-3’.

For generation of stable cell lines, EGFP and DFCP1/STX17 were cloned into the pLVX-IRES-Hygro vector (a gift from Harold MacGillavry) using the EcoRI and SpeI restriction sites. DFCP1 and STX17 were amplified by pcr using mCherry-DFCP1 (Itakura et al., 2012) and FLAG-STX17 (Kim et al., 2015) as templates. FLAG-STX17 was a gift from Noboru Mizushima (Addgene #45911) and mCherry-DFCP1 was a gift from Do-Hyung Kim (Addgene #86746).

### Cell culture and transfection

HeLa cells (ATCC) and HEK293T cells were cultured in Dulbecco’s modified Eagle’s medium containing 10% fetal calf serum and Penicillin/Streptomycin. Cells were maintained at 37 °C and 5% CO_2_ and were frequently tested for mycoplasma contamination using MycoAlert Mycoplasma Detection Kit (Lonza). Cells were plated on 25-mm diameter coverslips and transfected using Fugene6 transfection reagent (Roche) according to the manufacturer’s protocol. Experiments were started 1 day after transfection.

### Generation of stable cell lines

Cells stably expressing GFP-DFCP1 and GFP-STX17 were generated using a lentiviral expression system. Lentivirus packaging was performed by using MaxPEI-based co-transfection of HEK293T cells with psPAX2 (Addgene #12260), pMD2.G (Addgene, #12259) and the lentiviral vector pLVX-IRES-Hygro-EGFP-DFCP1 or pLVX-IRES-Hygro-EGFP-STX17. Supernatant of packaging cells was harvested up to 72 h of transfection, filtered through a 0.45-µm filter and incubated with a polyethylene glycol (PEG)-6000-based precipitation solution overnight at 4°C. After precipitation, virus was concentrated up to 100× by centrifugation and dissolution in 1× phosphate buffered saline (PBS). HeLa cells were incubated for 4 h in complete medium supplemented with 8 µg/ml polybrene before infection. To establish stable HeLa cell lines carrying EGFP tagged DFCP1 or STX17, medium was replaced 24–48 h after infection and 100 µg/ml Hygromycin (Invitrogen, ant-hg-5) was added.

Endogenous tagging of RAB7 was performed by CRISPR-Cas9 mediated genome editing (Ran et al., 2013). The HeLa GFP-RAB7 line was generated by transfecting HeLa cells (ATCC) with the donor plasmid pDONOR-GFP-RAB7 and pSpCas9(BB)-2A-Puro (PX459) V2.0 (Addgene, #62988) bearing the appropriate targeting sequence (5’-TAGTTTGAAGGATGACCTCT-3’; pX459v2-RAB7a sg3). 24 hours after transfection, Cas9-positive cells were selected by treatment with 1 µg/ml puromycin for 64 hours. Subsequently, cells were grown to confluency and seeded as single cells. EGFP-Rab7-KI was validated by PCR amplification using (FW; 5’-TACCCTGGCAAATGAGAGGC-3’ and RV; 5’-CTGGGCAAATGCTAGCGAAC-3’) and sequencing (5’-GGCTTAGCTCTAAGCCAATC-3’).Single clones were selected to have correct labeling of RAB7 by live cell microscopy and colocalization analysis with endo-lysosomal markers by immunofluorescence microscopy and were confirmed by sequence analysis.

### Fluorescence microscopy

Live-cell imaging was performed on a Spinning Disc Nikon Eclipse Ti Microscope with Perfect Focus System controlled by MetaMorph 7.7 software (Molecular Devices). An incubation chamber (Tokai Hit; INUBG2E-ZILCS) was used mounted on a motorized stage (ASI, MS-2000-XYZ). Coverslips were mounted in metal imaging rings immersed in medium. During imaging cells were maintained at 37°C and 5% CO2. Cells were imaged every 20 seconds for 1 hour using a 60x oil immersion objective (Plan Apo VC, N.A. 1.4, Nikon) and a Evolve 512 EMCCD camera (Photometrics). Vortran Stradus 445 nm, Cobolt Calypso 491 nm and Cobolt Jive 561 nm lasers were used for excitation. Aggregates were formed by addition of 500nM rapalog2 (Takara, B/B homodimerizer #635059) for 1 h, cells were imaged between 2-8hrs after aggregate formation.

For imaging of cells at specific timepoints, aggregates were induced and rapalog2 was washed out after 1 hr after which cells were imaged at 1h or 8 hr timepoints by taking 3 z-planes, 200 nm apart. Cells treated with Bafilomycin A1 were incubated with 200nM Bafilomycin A1 for 30 minutes before rapalog2 addition. Bafilomycin A1 was reapplied after rapalog2 washout.

### Image analysis

For analysis of different timepoints, first an average z-projection was made from the stack using ImageJ (NIH). An ROI was placed around the cells and ComDet v0.4.1 Plugin was used to detect particles using an approximated particle size of 4 pixels and a SNR of 8 for both mKeima channels. Data were transferred to Microsoft Excel, where the mKeima ratio per particle was calculated as the mKeima integrated intensity with 445 excitation divided by the total integrated intensity (at 445nm and 561nm excitation). Frequency distribution graphs were plotted using the GraphPad PRISM7 software. Graphs represent data from three independent experiments, with 10-15 cells per experiment. The percentage of cleared clusters is calculated as average percentage of the three independent experiments of clusters with an mKeima ratio <0.35 (sum from histogram frequencies with bin 0-0.3).

For analysis of individual cluster acidification, an ROI was placed around the particle and intensity was measured in both mKeima channels. Background intensities were measured in regions of the cell where no clusters were present. Data was transferred to Microsoft Excel and after background subtraction the mKeima ratio was calculated. Tracking of aggregates was performed by manual tracking using ImageJ.

The time interval between DFCP1 presence at the aggregate and acidification was calculated by taking the difference between the time at which the normalized DFCP1 signal dropped and stayed below 0.7 of the maximum intensity and the mKeima ratio stayed below 0.8.

For the averaged timing of aggregate acidification and STX17 presence, all the traced aggregates were aligned according to the first frame without STX17 presence as scored manually.

For analyzing average RAB7 intensity during aggregate acidification, individual events were aligned according to the frame at which mKeima ratio dropped halfway to the minimum value. Time points at which aggregates were out of focus or overlapped were not included in the analysis.

### FACS

FACS analysis was performed on a BD-Influx cell sorter. Measurements were made using a 488 (pH neutral) and 561 (pH low) nm laser with 630/22 nm and 610/20 nm emission filters, respectively. For each sample, 20,000 events were collected and subsequently gated for singlets and mKeima-positive cells. Data were analyzed using FlowJo v10 and plotted using MATLAB and GraphPad.

## Code availability

ComDet v.0.4.1 plugin for ImageJ is available online (https://github.com/ekatrukha/ComDet).

## ACKNOWLEDGMENTS

We would like to thank Harold MacGillavry for the pLVX-IRES-Hygro vector, Michael Davidson for the gift of mKeima-RED-N1, Noboru Mizushima for the FLAG-STX17, Do-Hyung Kim for mCherry-DFCP1, Didier Trono for the gift of pMD2.G and psPAX2 and Feng Zhang for pSpCas9(BB)-2A-Puro (PX459) V2.0. This research was supported by the European Research Council (ERC Consolidator Grant 819219 to L.K.) and the Dutch Research Council (NWO, ZonMW 91217002 to L. K.).

## AUTHOR CONTRIBUTIONS

A.J. and L.K. conceived research and designed the study. A.J. and G.K. created reagents, performed experiments, analysed data, and prepared the figures. W.N. helped with creating stable cell lines and E.K. helped with data analysis. A.J. and L.K. wrote the manuscript. L.K. supervised the study.

## FIGURES

**SUPPLEMENTARY FIGURE 1:**
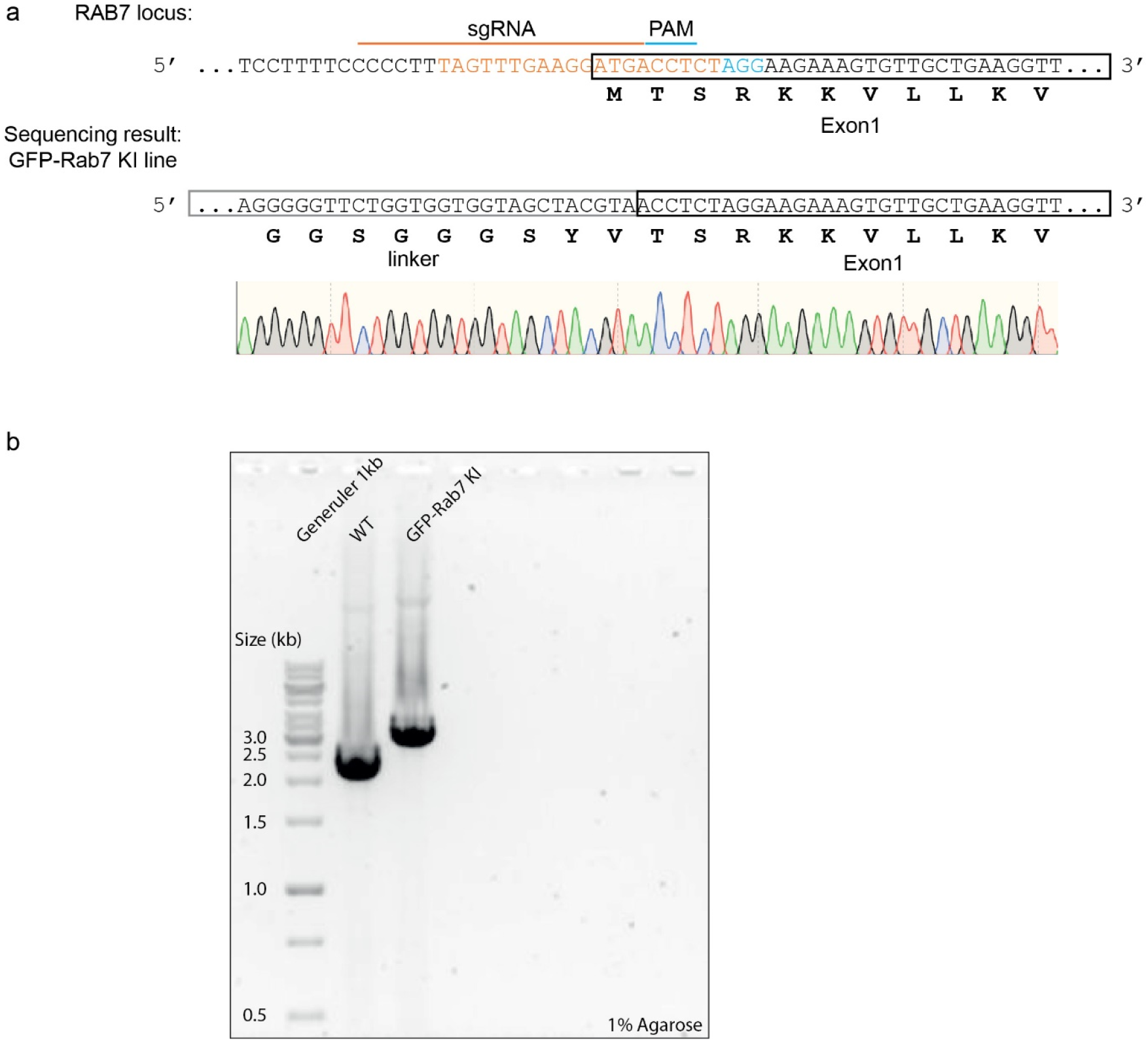
GFP-Rab7 KI HeLa line validation. a) Coding strand of knock-in target site before exon 1 (black box) of the *RAB7A* locus. The sgRNA target site (orange) and PAM sequence (cyan) are indicated. Insert is EGFP followed by a linker sequence (grey box). DNA sequencing of the homozygous knock-in clone is aligned below. b) 1% agarose gel including PCR-amplification product of *RAB7A* locus of *WT* and EGFP-Rab7 KI lines.

**SUPPLEMENTARY FIGURE 2:**
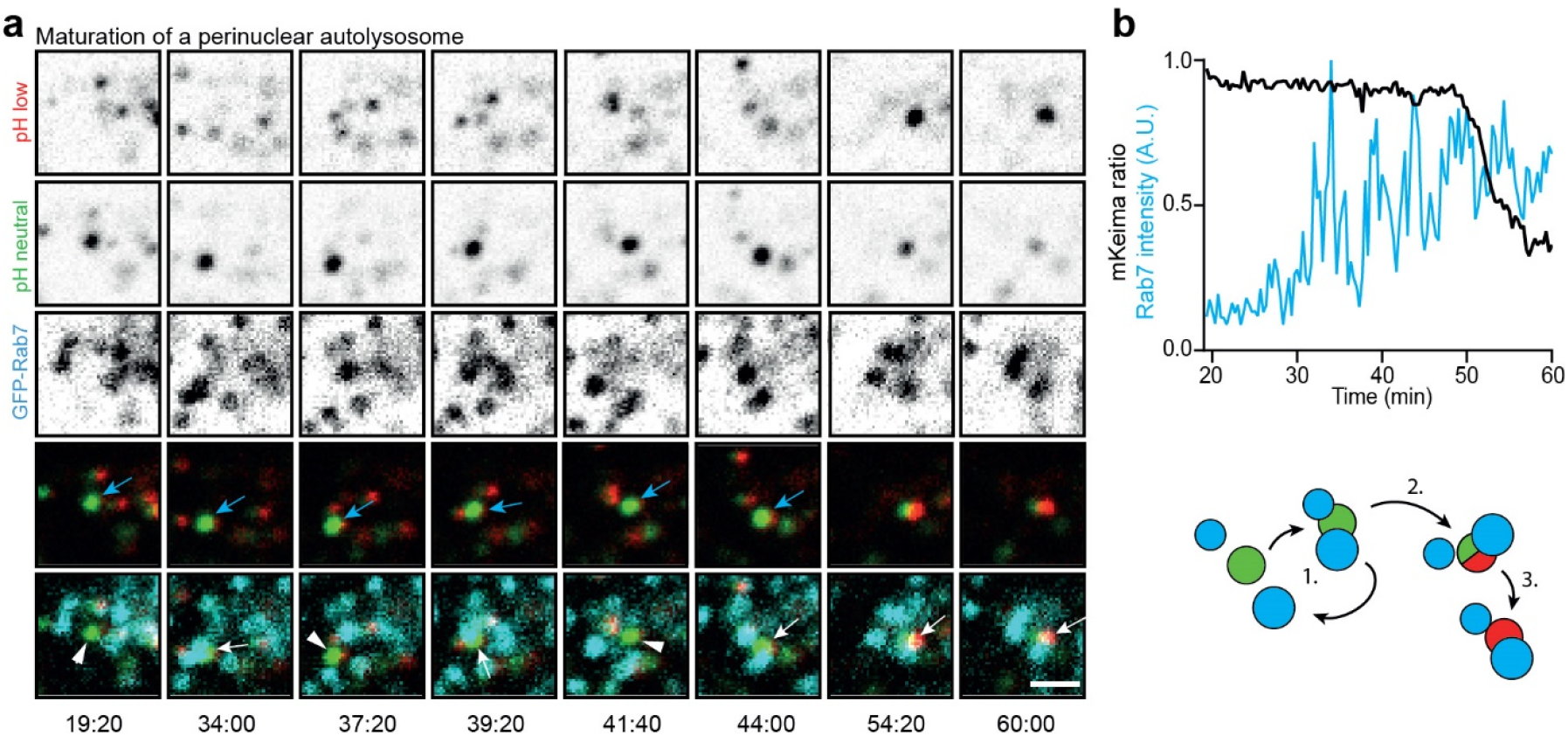
Maturation of a perinuclear autolysosome. a) Monitoring a forming autolysosome in the perinuclear region (see schematic in b). Frames from a time-lapse movie depicting multiple rounds of transient associations with RAB7 positive vesicles (1.), subsequent fusion (2.) and full acidification (3.). Arrows and arrowheads indicate interactions and absence of interactions, respectively. b) Analysis of RAB7 intensity and mKeima ratio of (a). Scale bar: 2 µm

### SUPPLEMENTARY VIDEO LEGENDS

**Video 1:** This video complements Figure 4c and shows transient STX17 (left) accumulation at mKeima-PIMs (middle) with concomitant acidification marked by green to red conversion of mKeima. Time denotes minutes:seconds. Scale bar marks 2 μm.

**Video 2:** This video complements Figure 6g and shows Rab7 (top left) accumulation at mKeima-PIMs (top right) with concomitant acidification marked by green to red conversion of mKeima. Time denotes minutes:seconds. Scale bar marks 2 μm.

**Video 3:** This video complements Supplemental Figure 2a and shows multiple rounds of transient associations of mKeima-PIM with RAB7 positive vesicles with concomitant gradual acidification. Time denotes minutes:seconds. Scale bar marks 2 μm.

